# Midgut barriers prevent the replication and dissemination of the yellow fever vaccine in *Aedes aegypti*

**DOI:** 10.1101/577213

**Authors:** Lucie Danet, Guillaume Beauclair, Michèle Berthet, Gonzalo Moratorio, Ségolène Gracias, Frédéric Tangy, Valérie Choumet, Nolwenn Jouvenet

## Abstract

**Background:** To be transmitted to vertebrate hosts *via* the saliva of their vectors, arthropod-borne viruses have to cross several barriers in the mosquito body, including the midgut infection and escape barriers. Yellow fever virus (YFV) belongs to the genus *Flavivirus*, which includes human viruses transmitted by *Aedes* mosquitoes, such as Dengue and Zika viruses. The live-attenuated YFV-17D vaccine has been used safely and efficiently on a large scale since the end of World War II. Early studies have shown, using viral titration from salivary glands of infected mosquitoes, that YFV-17D can infect *Aedes aegypti* midgut, but does not disseminate to other tissues.

**Methodology/Principal Findings:** Here, we re-visited this issue using a panel of techniques, such as RT-qPCR, Western blot, immunofluorescence and titration assays. We showed that YFV-17D replication was not efficient in *Aedes aegypti* midgut, as compared to the clinical isolate YFV-Dakar. Viruses that replicated in the midgut failed to disseminate to secondary organs. When injected into the thorax of mosquitoes, viruses succeeded in replicating into midgut-associated tissues, suggesting that, during natural infection, the block for YFV-17D replication occurs at the basal membrane of the midgut. Our NGS analysis revealed that YFV-Dakar genome exhibited a greater diversity than the vaccine strain; a trait that may contribute to its ability to infect and disseminate efficiently in *Ae. aegypti*.

**Conclusions/Significance:** The two barriers associated with *Ae. aegypti* midgut prevent YFV-17D replication. Our study contributes to our basic understanding of vector–pathogen interactions and may also aid in the development of non-transmissible live virus vaccines.

**Author summary:** Most flaviviruses, including yellow fever virus (YFV), are transmitted between hosts by mosquito bites. The yellow fever vaccine (YFV-17D) is one of the safest and most effective live virus vaccine ever developed. It is also used as a platform for engineering vaccines against other health-threatening flaviviruses, such as Japanese encephalitis, West Nile, Dengue and Zika viruses. We studied here the replication and dissemination of YFV-17D in mosquitoes. Our data showed that YFV-17D replicates poorly in mosquito midgut and is unable to disseminate to secondary organs, as compare to a YFV clinical isolate. Our study contributes to our basic understanding of the interactions between viruses and their vectors, which is key for conceiving new approaches in inhibiting virus transmission and designing non-transmissible live virus vaccines.

## Introduction

Arborviruses, which are transmitted among vertebrate hosts by blood-feeding arthropod vectors, put billions of people at risk worldwide. Viral infection in arthropod is usually persistent. Following uptake of an infectious blood meal by a female mosquito, arbovirus must initiate a productive infection of the midgut epithelium, which consists of a single layer of cells [1]. To develop a disseminated infection, virus must then escape the midgut into the haemocoel and infect secondary tissues such as the fat body, trachea and the salivary glands [1]. Finaly, the virus needs to be released into salivary ducts for horizontal transmission to an uninfected vertebrate host [1]. Traditional means of controlling the spread of arbovirus infection include mosquito control and vaccination of susceptible vertebrates. However, in many cases, these measures are either unavailable or ineffective. To successfully implement the strategy of blocking the virus at the arthropod stage, further knowledge of the virus/vector interactions is required.

Flaviviruses constitute the most important and diverse group of arthropod-transmitted viruses causing diseases in humans. They are 50 nm-diameter enveloped viruses harboring a single positive-strand RNA genome of 10.7 kb. The genome encodes a polyprotein that is cleaved into seven non-structural (NS) proteins (NS1, NS2A, NS2B, NS3, NS4A, NS4B, and NS5) and three structural proteins: capsid (C), pre-membrane/membrane (prM/M) and envelope (Env). The C, M, and Env proteins are incorporated into virions, while NS proteins are not [2, 3]. NS proteins coordinate RNA replication, viral assembly and modulate innate immune responses.

Several members of the flavivirus genus, such as dengue virus (DENV), yellow fever virus (YFV) and Zika virus (ZIKV) are highly pathogenic to humans and constitute major global health problems. YFV is responsible for viral hemorrhagic fever resulting in up to 50% fatality [4]. Despite the existence of the safe and effective live-attenuated vaccine YFV-17D, YFV regularly resurges in the African and South American continents, as illustrated by recent outbreaks in Brazil and equatorial Africa [5–7]. The YFV-17D vaccine has been used safely and efficiently on a large scale since the end of World War II [8]. It was developed in the 1930’s by passaging the blood of a human patient in rhesus macaques and later in mouse and chicken embryo tissues [9]. A single dose confers protective immunity for up to 35 years. During the attenuation process, YFV-17D has lost its neurotropic and viscerotropic properties, which account for the major disease manifestations of yellow fever in primates [10, 11]. The molecular determinants responsible for its virulence attenuation and immunogenicity are poorly understood. We have recently shown that YFV-17D binds and enters mammalian cells more efficiently than a non-attenuated strain, resulting in a higher uptake of viral RNA into the cytoplasm and consequently a greater cytokine-mediated antiviral response [12]. This differential entry process may contribute to attenuation in humans.

YFV-17D is also used as a platform for engineering vaccines against other health-threatening flaviviruses, such as vaccines against Japanese encephalitis virus (JEV), West Nile virus (WNV), the four serotypes of DENV, and, more recently, ZIKV [13–16]. These vaccines consist in a YFV-17D backbone in which sequences coding for prM/E proteins are replaced by those of the selected flavivirus. Some of these live-attenuated chimeric vaccines are commercially available [17, 18], with variable success [19]. YFV-17D is thus a key component in controlling flaviviral disease and it must not disseminate in mosquitoes. Early studies have shown, using almost exclusively viral titration by plaque assays, that YFV-17D can infect *Aedes aegypti* midgut, [20, 21], but does not disseminate to other tissues and fails to be transmitted to a novel host. Here, we re-visited this question using a variety of techniques and showed that not only the midgut escape barrier, but also the midgut infection barrier, restrict YFV-17D replication in its vector.

## Materials and methods

### Viruses

The YFV-17D vaccine strain (YF-17D-204 STAMARIL, Sanofi Pasteur, Lyon) was provided by the Institut Pasteur Medical Center. The YFV-DAK strain (YFV-Dakar HD1279) was provided by the World Reference Center for Emerging Viruses and Arboviruses (WRCEVA), through the University of Texas Medical Branch at Galveston, USA. Viral stocks were prepared on Vero cells, concentrated by polyethylene glycol 6000 (Sigma) precipitation and titrated on Vero cells by plaque assay as described previously [22].

### Cells

The Aag2 mosquito cell lines (provided by the teams of M. Flamand and L. Lambrechts, Institut Pasteur, Paris) are derived from larvae of *Aedes aegypti*. They were cultured in a humid chamber at 28°C, with no CO_2_, in Leibovitz medium (Gibco™ Leibovitz’s L-15 Medium, Life Technologies) supplemented with 10% fetal bovine serum (FBS), 2% tryptose phosphate buffer (Gibco™ Tryptose Phosphate Broth 1X, Life Technologies), 1% non-essential amino acid solution (Gibco™ NEAA 100X MEM, Life Technologies), 1% penicillin-streptomycin (P/S) (Sigma). Vero cells, which are African green monkey kidney epithelial cells, were purchased from the American Type Culture Collection (ATCC) and used to perform viral titration. They were maintained in Dulbecco’s modified Eagle’s medium (DMEM, Invitrogen), supplemented with 10% FBS and 1% P/S.

### Antibodies

Env MAb 4G2 hybridoma cells were kindly provided from P. Desprès (La Réunion University, Sainte Clotilde). Anti-YFV-NS4B and anti-DENV NS1 antibodies (that recognize YFV-NS1), were kind gifts from C.M. Rice (Rockefeller University, NY) [23] and M. Flamand (Institut Pasteur, Paris) [24], respectively. Anti-actin (A1978, Sigma) and anti-tubulin (T5168, Sigma) antibodies were used as loading controls for mosquito organs and Aag2 cells, respectively. Secondary antibodies were as followed: anti-mouse 680 (LI-COR Bioscience), anti-rabbit 800 (Thermo Fisher Scientific) and anti-rabbit Cy3 (Life Technologies).

### Infection and dissection of mosquito

The Paea strain of *Ae. aegypti* is a laboratory colony originated from mosquitoes collected in French Polynesia in 1960 and conserved in the laboratory since 400–450 generations. Adult mosquitoes were maintained at 25 ± 1 °C and 80% relative humidity with a light/dark ratio of 12 h/12 h. The larvae were provided with brewer’s yeast tablets and adults were given continuous access to 10% sucrose solution. Sucrose was removed 24 h prior to the infectious blood meal. The infectious blood meal was comprised of half-human blood and half-viral suspension (4.10^7^ PFU/mL in the mix). The blood donors were randomly selected from a population of healthy volunteers donating blood at the ‘Etablissement Français du Sang’ (EFS), within the framework of an agreement with Institut Pasteur. Experimental procedures with human blood have been approved by EFS Ethical Committees for human research. All samples were collected in accordance with EU standards and national laws. Informed consent was obtained from all donors. Seven day-old female mosquitoes were allowed to feed for 15 min through a collagen membrane covering electric feeders maintained at 37°C (Hemotek system). Blood-fed females were selected and transferred into cardboard boxes protected with mosquito nets. Alternatively, ice-chilled mosquitos were injected intrathoracically with twice 69 nL of viral stock with a micro-injector (Drummond, Nanoject II). Mosquitoes were anesthetized on ice at various time-points after infection. They were passed through a 70% ethanol bath and then in a PBS bath before being dissected in a drop of PBS under a magnifying glass using tweezers. The midguts, legs and salivary glands were removed and placed in a tube containing sterilized glass beads of a diameter of 0.5 mm (Dutscher) in a suitable lysis buffer. Experiments were reproduced in triplicate with 5–10 mosquitoes collected at each time-point for dissection.

### RT-qPCR analysis

The mosquito organs were crushed using a tissue homogenizer (Ozyme, Precellys Evolution) during twice 15 s at 10000 g. Total RNA was extracted from mosquito tissues with the NucleoSpin RNA II kit (Macherey-Nagel). YFV RNA was quantified using NS3-specific primers and TaqMan probe (NS3-For CACGGCATGGTTCCTTCCA; NS3-MFAM CAGAGCTGCAAATGTC; NS3-Rev ACTCTTTCCAGCCTTACGCAAA) with TaqMan^®^ RNA-to-CT™ 1-Step (Thermo Fisher Scientific) on a QuantStudio 6 Flex machin (Applied Biosystems). Genome equivalent (GE) concentrations were determined by extrapolation from a standard curve generated from serial dilutions of total YFV RNA of known concentration.

### Western blot analysis

Individual midguts and salivary glands were collected in RIPA buffer (Sigma) containing protease inhibitors (Roche Applied Science). Tissue lysates were normalized for protein content with Pierce 660nm Protein Assay (Thermo Scientific), boiled in NuPAGE LDS sample buffer (Thermo Fisher Scientific) in non-reducing conditions and 32 µg (midgut) or 14 µg (salivary glands) of proteins (corresponding to around 10 pooled organs) were separated by SDS-PAGE (NuPAGE 4-12% Bis-Tris Gel, Life Technologies). Separated proteins were transferred to a nitrocellulose membrane (Bio-Rad). After blocking with PBS-Tween-20 0,1% (PBST) containing 5% milk for 1 hour at RT, the membrane was incubated overnight at 4°C with primary antibodies diluted in blocking buffer. Finally, the membranes were incubated for 1 hour at RT with secondary antibodies diluted in blocking buffer, washed, and scanned using an Odyssey CLx infrared imaging system (LI-COR Bioscience).

### Immunofluorescence

After dissection, individual midgut were deposited on slides, fixed in cold acetone for 15 min and rehydrated in PBS for 15 min. The midguts were then incubated for 2 h in Triton X-100 (0.2%). After washing with PBS, they were incubated for 30 min with PBS + 0.1% Tween 20 + 1% BSA. The slides were then incubated overnight at 4°C with anti-YFV-NS4B antibodies diluted 1:1000 in PBS. After washing with PBS, they were incubated for 1 h with secondary antibodies and washed with PBS. The actin network was visualized with phalloidin Alexafluor 488 (Invitrogen). After washing, nuclei were stained using Prolong gold antifade containing 4′,6-diamidino-2-phenylindole (DAPI) (Invitrogen). All preparations were observed with a confocal microscope (ZEISS LSM 700 inverted) and images were acquired with the ZEN software.

### Deep-sequencing of viral stocks

Viral RNAs were extracted from 20 µl of viral stock using Trizol (Ambion, TRIzol ™ Reagent), were re-suspended in 20 µL of RNAse-free water and treated with DNAse with the DNA-free kit (Ambion) before being stored at -80°C. Synthesis of cDNAs was carried out with the Maxima H Minus First Strand kit (Thermo Fisher Scientific) from 250 ng of viral RNA. Three fragments of the viral genome were amplified by PCR using the Phusion^®^ High-Fidelity DNA Polymerase kit (NEB) using primers mainly described previously [25]. New primers targeting the 3’-UTR of the genome were designed for optimal amplification of YFV-17D and YFV-DAK (Table 1). The PCR products were purified with the NucleoSpin^®^ Gel kit and PCR Clean Up (Macherey-Nagel), resuspended in 40 µL of RNAse-free water and stored at -20°C. The PCR products were fragmented randomly with the NEBNext^®^ dsDNA fragmentase kit (NEB) and then purified with the AMPure^®^ XP Beads kit (Beckman Coulter, Inc.). The Illumina sequencing library were prepared with the NEBNext^®^ Ultra DNA Library Prep kit (NEB) by selecting 400-bp fragments. NEBNext^®^ Multiplex Oligos for Illumina^®^ primers (NEB) were used. Purification was performed with the AMPure^®^ XP Beads kit and quantification using the Qubit ™ dsDNA BR Assay kit (Thermo Fisher Scientific). Samples from the library, diluted to 4 nM, were sequenced on a NextSeq^®^ 500 sequencer (Illumina) machine with the NextSeq^®^ 500 Mid Output Kit v2 kit (150 cycles) (Illumina), to generate read of 150 bp.

**Table 1.**
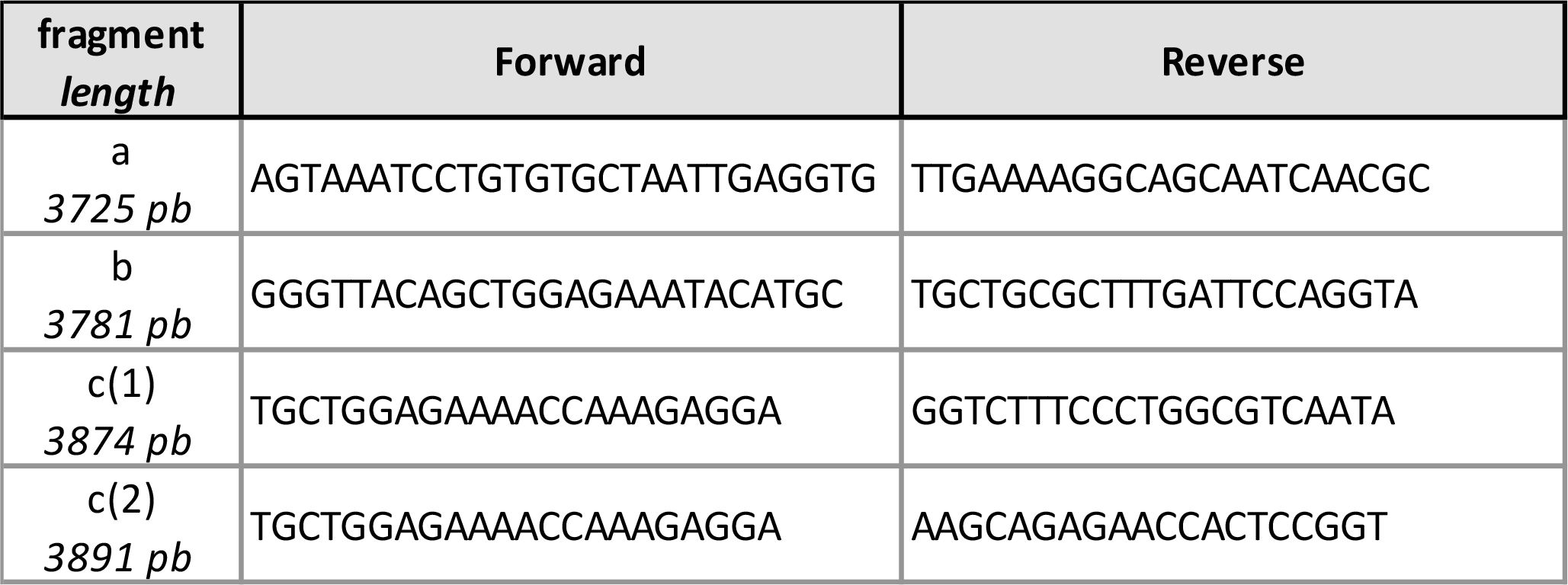
Primers used for YFV sequencing. Viral genome was amplified in three fragments (a, b and c) of approximately 3000 pb. Different reverse primers were used to amplify the c fragment of the two YFV strains resulting in c(1) for YFV-17D and c(2) for YFV-Dakar. All primers used to amplify fragments a and b were previously desccribed [25].

### Viral sequence analysis and comparison of diversity

Reads were trimmed for adapters and low quality ones were filtered using Trim Galore! (www.bioinformatics.babraham.ac.uk/projects/trim_galore/) using the following parameters: quality 30, length 100 and stringency 4. Final reads quality was evaluated using FastQC (www.bioinformatics.babraham.ac.uk/projects/fastqc/). Reads were aligned on the YFV-Asibi reference genome AY640589.1 using BWA [26] and SAMtools [27]. PCR duplicates were removed from alignment using Picard Tools MarkDuplicates (http://broadinstitute.github.io/picard/). Consensus sequence was obtained using SAMtools mpileup, VarScan mpileup2cns (min-var-freq 0.5) and BCFtools consensus [27]. For YFV-DAK, previous steps were repeated to solve drop of coverage issues. New alignments were made against respective consensus sequences. Variant determination was done with VarScan mpileup2snp (min-var-freq 0.01, strand-filter 0). The diversity index ’Shannon entropy’ was calculated using DiversiTools (btcutils) (http://josephhughes.github.io/DiversiTools/), as described previously [28]. Simpson’s diversity index (1-D) was calculated using a method previously described [29]. Graphs were generated with R script and R Studio programs [30], ggplot2 [31], readr (https://cran.r-project.org/web/packages/readr/index.html), Seqinr and gridExtra (https://cran.r-project.org/web/packages/gridExtra/index.html) packages.

### Statistical analysis

Data were analyzed using GraphPad Prism 7. Statistical analyses were performed with multiple t tests, Wilcoxon test or Mann-Whitney test (* p < 0.05 ; ** p < 0.01 ; *** p < 0.001 ; **** p < 0.0001, ns, not significant.), as indicated.

## Results

### YFV-17D, but not YFV-DAK, fails to overcome the midgut barriers of *Aedes aegypti*

The replication and dissemination of YFV-17D was studied in the *Ae. aegypti* strain Paea. The clinical isolate YFV-Dakar HD1279 (YFV-DAK), whose replication in Rhesus Macaque is well characterised [32], was used as a positive control for these experiments. Virus produced on Vero cells were mixed with human blood to prepare a meal containing 4.10^7^ PFU/mL of either YFV-17D or YFV-DAK. Five to ten mosquitoes were collected every 2-3 days until 14 days post-feeding (dpf). Mosquitoes were dissected to separate the midgut from legs and salivary glands. Virus production in these tissues was first assayed by calculating the viral titer by plaque assays on Vero cells. Several whole mosquitoes were also analyzed 20 minutes after feeding to ensure that the mosquitoes ingested a similar amount of viral particles of both viral strains. Around 10^3^ infectious particles of YFV-DAK were detected per midguts 3 dpf (Fig. 1A). Viral titers remained high in midguts until 14 dpf. YFV-DAK infectious particles were present in legs as early as 5 dpf and in salivary glands as early as 7 dpf (Fig. 1A). This replication pattern is comparable to that of South American and Africain YFV isolates in the strain *Ae. aegypti* AE-GOI [33]. Midgut of mosquitoes infection with YFV-17D produced 1 to 2 log less infectious particles than YFV-DAK at 3 dpf (Fig. 1B). Infectious particles were detected in a unique leg sample at 14 dpf. No virus was detected in salivary glands of mosquitoes infected with YFV-17D. Thus, by contrast to YFV-DAK, and in agreement with previous studies performed with the *Ae*. *aegypti* strains Rexville or Rexville-D (Rex-D) [21, 34-36], YFV-17D disseminated poorly in the strain Paea.

**Figure 1.**
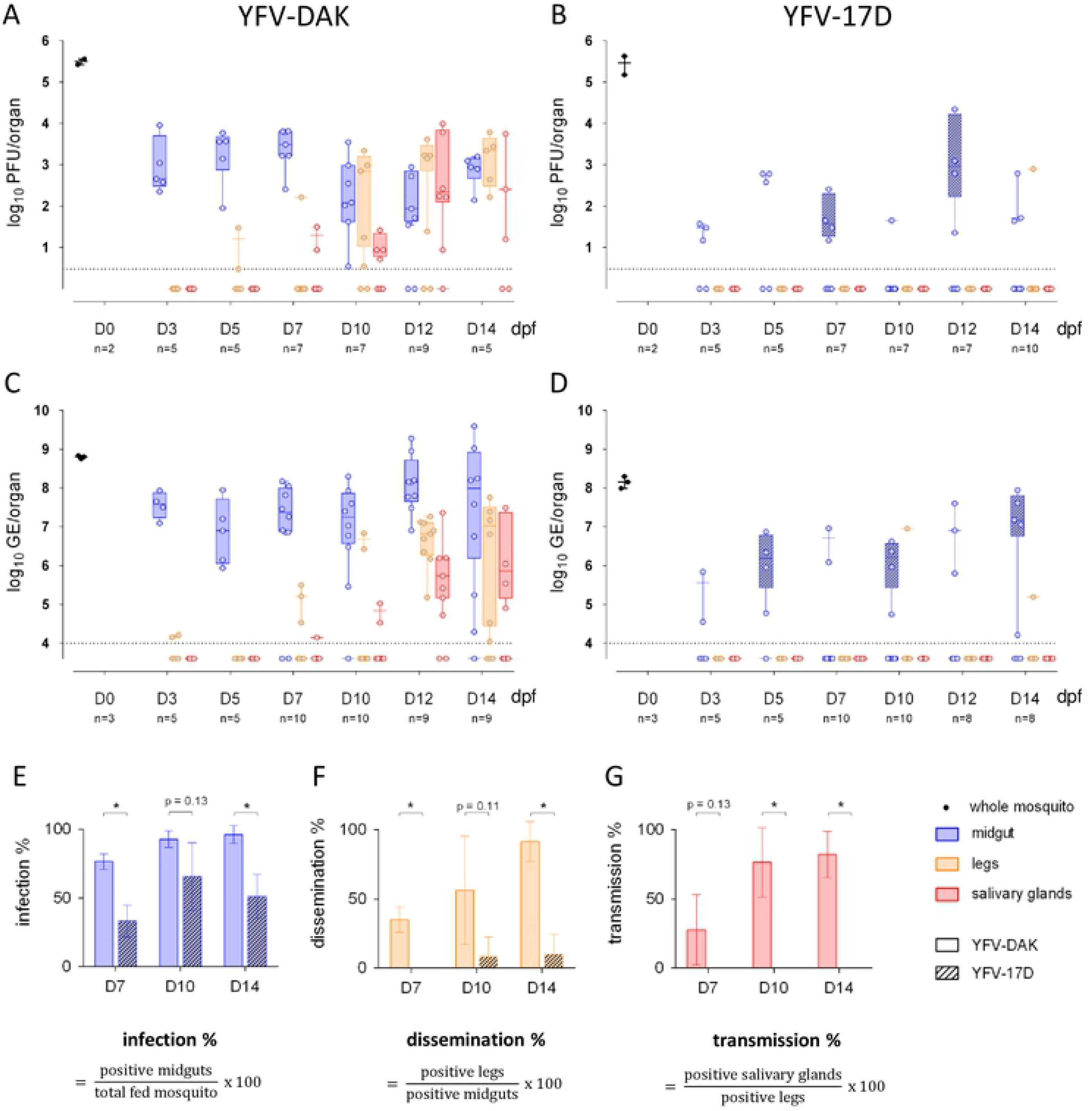
YFV-17D, but not YFV-DAK, fails to overcome the midgut barriers of *Aedes aegypti*. Mosquitoes were orally infected with 4.10^7^ PFU/mL of YFV-DAK (A and C) or YFV-17D (B and D). (A and B) The presence of infectious viruses in individual midgut, legs and salivary glands was assessed by plaque assay on Vero cells at 3, 5, 7, 10, 12 and 14 day post feeding (dpf). Several whole mosquitoes were also analyzed 20 minutes after feeding (black dots). Each data point represents the YFV titers of a single organ. (C, D) The relative amounts of organ-associated viral RNA were determined by RT-qPCR analysis and are expressed as genome equivalents (GE) per organ at 3, 5, 7, 10, 12 and 14 dpf. Total RNA was also extracted from several whole mosquitoes the same day of the feeding (black dots). (A-D) The dashed lines indicate the limit of detection. Data shown were obtained from one representative experiment. YFV infection rates among midguts (E), YFV dissemination rate among legs (F) and YFV transmission rate among salivary glands (G) were determined by RT-qPCR analysis at 7, 10 and 14 dpf. (E-G) Data were obtained from 3 independent experiments. Error bars indicate the means ± SD. Statistical analyses were performed using multiple t tests (* p < 0.05).

Viral replication was assessed in the midguts, legs and salivary glands by measuring viral RNA quantity over-time by RT-qPCR. Total RNA was also extracted from several whole mosquitoes the same day of the feeding to insure that they had ingested similar amount of infectious particles from both viral strains. Around 10^7^ copies of viral RNA were detected in midguts of mosquitoes infected with YFV-DAK since 3 days (Fig. 1C). The viral RNA copy number per midgut remained high until 14 dpf, indicating that viral replication had already reached a plateau at early stage of infection (Fig. 1C). In agreement with titration assays (Fig. 1A), YFV-DAK RNA was detected in legs and salivary glands of mosquitoes around 7 dpf. The quantity of viral RNA detected in these secondary organs increased over time to reach on average 10^7^ copies RNA in legs and 10^6^ copies in salivary glands at 14 dpf (Fig. 1C). Around 5.10^5^ copies of YFV-17D RNA was detected in 2 out of 5 midguts of blood-feed mosquitoes at 3 dpf (Fig. 1D). At 12 dpf, around 10^7^ copies of YFV-17D RNA was detected in 4 out of 8 mosquitoes, which is 10 time less than in YFV-DAK infected moquitoes. YFV-17D RNA was found in legs of 2 mosquitoes among the 46 blood-fed mosquitoes collected during 14 days. No virus was dectected in the salivary glands of these 46 mosquitoes (Fig. 1D). Thus, in agreement with our titration assays (Fig. 1B) and with previous studies performed with Rexville strains of *Ae*. *aegypti* [34–36], YFV-17D disseminated poorly in the strain Paea. The RT-qPCR analysis also revealed that the vaccine strain replicated less efficiently than YFV-DAK in its vector.

The pourcentage of mosquitoes that were positive for viral RNA among the mosquitoes that had taken blood was calculated based on RT-qPCR data obtained from 3 independent experiments (Fig. 1E). Significantly less midguts were postivive for YFV-17D RNA than YFV-DAK RNA at days 7 and 14 post-feeding, suggesting that the midgut infection barrier restrict the replication of the vaccine strain. Viral dissemination was defined by the presence of viral RNA in the legs of mosquitoes whose midguts were infected (Fig. 1F). YFV-DAK had disseminated in around 40% of infected mosquitoes at 7 dpf and in around 90% of mosquitoes at 14 dpf. At this time, YFV-17D had disseminated in around 10% of them (Fig. 1F). YFV-DAK dissemination rates are consistent with the ones reported for the YFV-Asibi strain [36] or clinical isolates from Peru [35] in Rexville mosquitoes. YFV-DAK RNA was detected in salivary glands of approximately 75% of mosquitoes whose legs were infected, revealing that the virus was efficiently transmitted (Fig. 1G). By constrast, no viral tranmission was observed for YFV-17D.

To investigate the replication ability of the two viral strains further, the presence of viral antigens in pooled midguts and salivary glands of mosquitoes fed on blood containing 4.10^7^ PFU/mL of either YFV-17D or YFV-DAK was analyzed by Western blots at days 7 and 14 post-feeding using antibodies against Env and NS1. The Env protein was detected at both time-points in the midgut of mosquitoes infected with the strain YFV-DAK, in a majority form of around 45 kDa and a minor form of around 35 kDa (Fig. 2A). The Env protein was not detected in the salivary glands 7 days after the blood meal but was present as a 45 kDa form 14 days after the blood meal (Fig. 2A). These data are in good agreement with the titration and RT-qPCR data presented in Figure 1. Like the Env protein, the NS1 protein was detected in the midguts of mosquitoes infected with the YFV-DAK strain at both 7 and 14 dpf (Fig. 2A). In midguts, NS1 was detected at the expected size of 45 kDa, but also as heavier forms of around 80 kDa. These forms could represent NS1-2A, a polyprotein precursor consisting of NS1 and a portion of NS2A. This NS1-2A form was previously reported in human SW-13 cells infected with YFV-17D [23] and maybe generated by alternative cleavage sites in the NS2A region upstream from the cleavage site generating the N-terminus of NS2B. Alternatively, they could represent glycosylated versions of NS1 monomer or dimer. NS1 was also detected in the salivary glands of mosquitoes infected with YFV-DAK for 14 days (Fig. 2A). No or very little signal was detected by the anti-NS1 or anti-Env antibodies in organs of mosquitoes infected with YFV-17D (Fig. 2A). In order to ensure that the antibodies directed against the NS1 and Env proteins recognize YFV-17D proteins, control experiments were performed with the *Ae. aegypti* Aag2 cells infected for 24 or 48 hours at an MOI of 0.1 with both viral strains. Both proteins were well detected in cells infected for 48 hrs, independently of the viral strain used (Fig. 2B). Thus, absence of detection of YFV-17D Env and NS1 proteins in the mosquito organs at 7 and 14 dpf is not due to poor recognition of the viral antigens, nor the antibodies used, but reflects a low-level replication. These data confirm our titration and RT-qPCR analysis (Fig. 1). Of note, the YFV-DAK Env was detected as 2 forms in Aag2 cells infected for 48 hours while the YFV-17D Env was detected as a unique form. No YFV-17D proteins were detected at 24 hours post-infection, suggesting that the replication of the vaccine strain is less efficient in Aag2 cells than the one of YFV-DAK.

**Figure 2.**
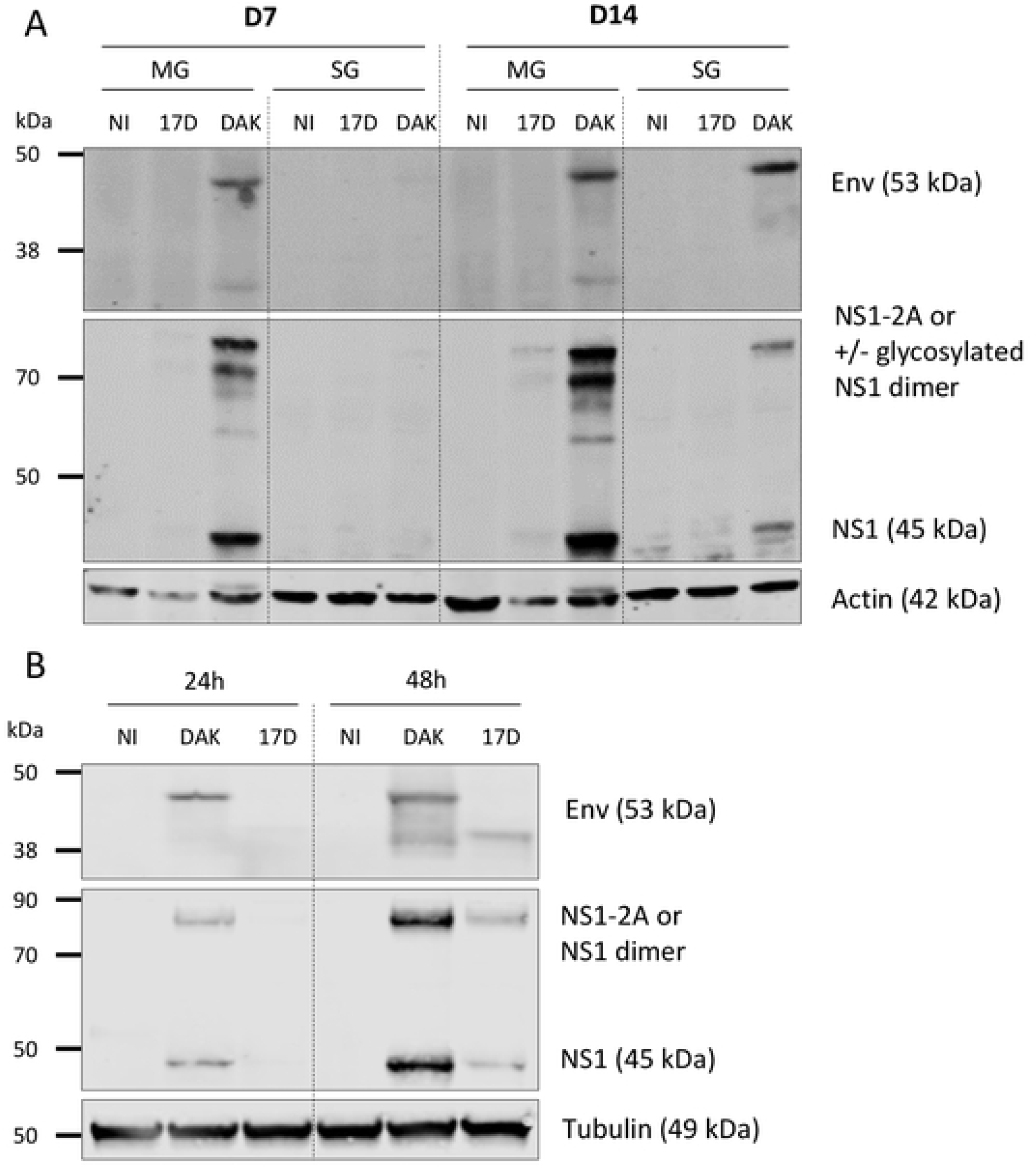
YFV-DAK produces detectable levels of viral proteins in infected mosquitoes. (A) Mosquitoes were fed with human blood containing 4.10^7^ PFU/mL of YFV-DAK or YFV-17D or with no virus (NI). The presence of viral antigens in 10 pooled midguts and salivary glands was analyzed by immunoblotting at 7 and 14 dpf using antibodies recognizing actin, viral NS1 or Env proteins. (B) Aag2 cells were carried infected at a MOI of 1. Whole-cell lysates were analyzed by immunoblotting at the indicated times post-infection using antibodies recognizing human tubulin, viral NS1 or Env proteins. Non-reducing condition were used to detect the Env proteins.

Finally, to confirm RT-qPCR and Western blot data, immunofluorescence analyses were performed on miguts of mosquitoes fed since 7 days using antibodies againt the viral protein NS4B. YFV-DAK antigens were evenly distributed in foci over the entire epithelium at this time (Fig. 3A). By contrast, YFV-17D antigens were found in one or two localized foci in infected midguts. In an attempt to assess this unevenly distribution of YFV-17D replication sites, the midgut of mosquitoes infected with both viral strains for 3 or 7 days were cut longitudinally into two equal parts. The presence of viral RNA was determined by RT-qPCR analysis performed on individual half midguts (Fig. 3B). Among 18 mosquitoes that injested blood containing YFV-17D, 3 half midguts were positive for YFV-17D RNA at day 3 post-infection and only 2 at day 7 post-infection. This is in agreement for our previous results (Fig. 1E). Among these five positive midguts, only one contained YFV-17D RNA in both sections (Fig. 3B). As expected based on previous results (Fig. 1E), it was easier to obtain midguts positive for YFV-DAK. Twelve out of the 15 midguts that were positive for YFV-DAK RNA contained viral RNA in both sections. These experiments revealed that YFV-17D replication in *Ae. aegypti* midgut is more confined that the one of YFV-DAK.

**Figure 3.**
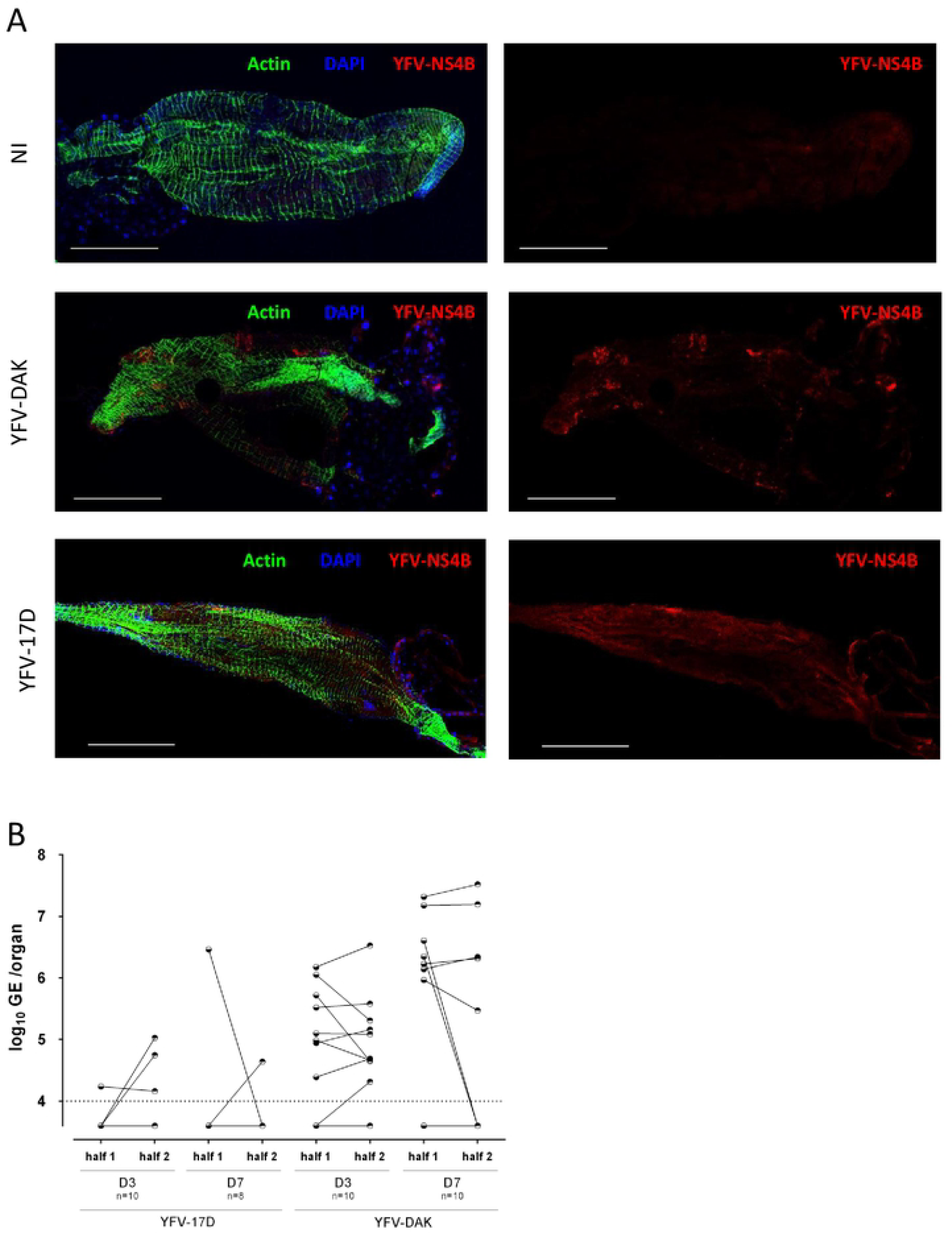
Replication of YFV-17D in midguts is localized to confined area. Mosquitoes were fed with human blood containing 4.10^7^ PFU/mL of YFV-DAK, YFV-17D or with no virus (NI). (A) Midguts were dissected at 7 dpf and stained with DAPI to visualize nuclei (blue), Phalloidin Texas Red to visualize actin (green) and with antibodies specific the viral protein NS4B (red). Images were acquired with confocal microscope equipped with a x40 objective. Scale bars are 0,5 mm. (B) The midguts of infected mosquitoes were cut longitudinally into two parts at 3 or 7 dpf. The presence of viral RNA was determined by RT-qPCR analysis performed on individual half midguts. The data are expressed as genome equivalents (GE) per organ. The dashed line indicates the limit of detection.

Together, these data show that, by contrast the clinical isolate YFV-DAK, the vaccine strain replicated poorly in, and disseminated poorly from *Ae. aegypti* midgut.

### YFV-17D and YFV-DAK replicate in the midgut when mosquitoes are inoculated intra-thoracically

To assess whether YFV-17D could infect *Ae. aegypti* when delivered *via* a non-oral route, mosquitoes were inoculated intra-thoracically with 2,5.10^4^ PFU of YFV-17D or YFV-DAK. This amount of infectious particles injected corresponds to around 10 times less PFU than when mosquitoes are taking around 5 µL blood meal containing 4.10^7^ PFU/mL. The presence of viral RNA was analyzed by RT-qPCR 10 days after injection. Mosquitoes infected *via* a blood meal served as controls. Several whole mosquitoes were also analyzed 20 minutes after feeding or injection to ensure that a similar amount of viral particles of both viral strains were delivered in mosquitoes. In good agreement with our previous experiments (Fig. 1), around 35% of midguts (8 out 22) were positive for YFV-17D RNA, whereas 81% (18 out 22) were positive for YFV-DAK RNA 10 dpf (Fig. 4A). Moreover, significantly less viral RNA (around 10 times) was found in YFV-17D-infected midguts as compared to DAK-infected midguts (Fig. 4A). YFV-DAK RNA was detected in legs and salivary glands of around 50% of these mosquitoes. By contrast, YFV-17D was detected in the legs of a unique mosquito out of 22 and was not detected in salivary glands (Fig. 4A), confirming the inability of the vaccine strain to spread to secondary organs and to be transmitted when orally delivered. When the midgut barriers were bypassed by injecting *Ae. aegypti* mosquitoes in the thorax, 100% of midguts were positive for both viral strains and similar amounts of YFV-17D and YFV-DAK RNA were detected in this organ, indicating that both viral strains successfully replicated in midgut associated tissue when bypassing the lumen (Fig. 4B). All legs and salivary glands are positive for YFV-17D and YFV-DAK RNA (Fig. 4B), revealing that the two viruses were efficiently infecting secondary tissues once the midgut was bypassed. Of note, significantly more (around 10 times) YFV-DAK RNA was detected in salivary glands than YFV-17D RNA (Fig. 4B), suggesting that YFV-17D is sensitive to the salivary gland infection barrier.

**Figure 4.**
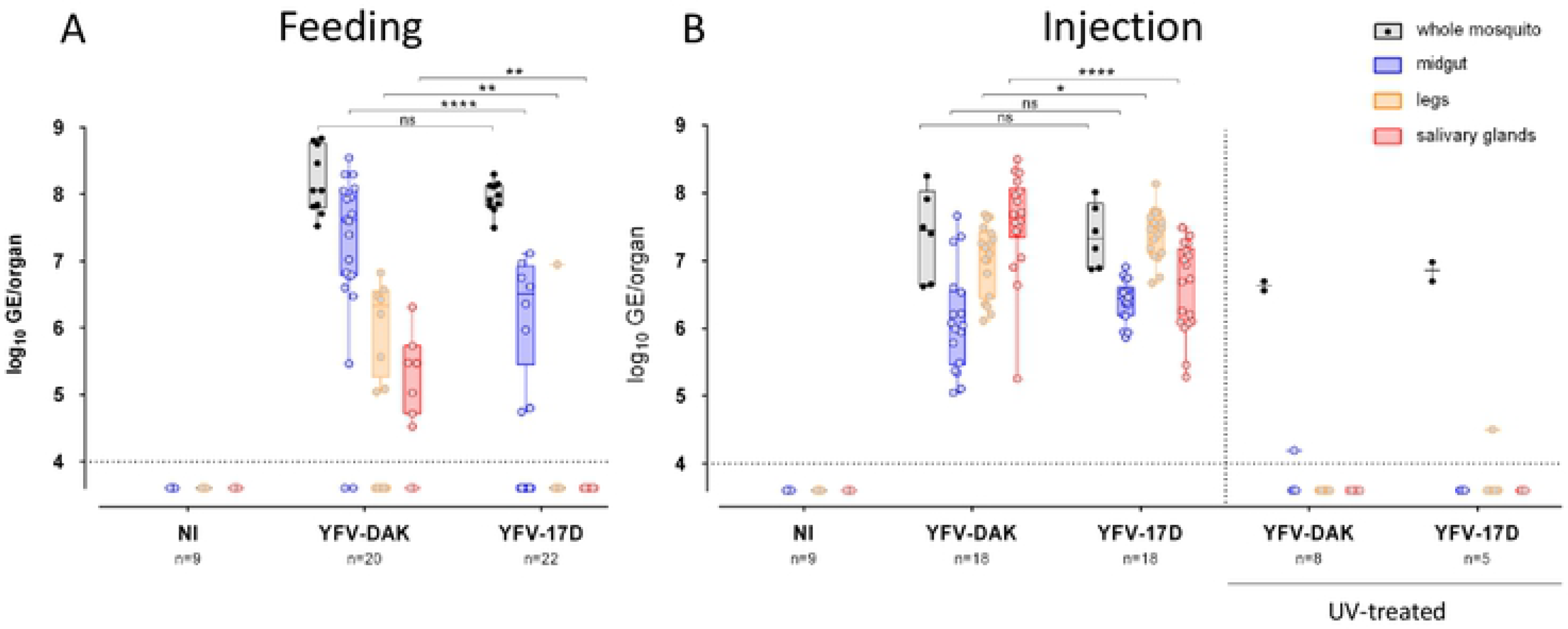
YFV-17D and YFV-DAK replicate in secondary organs when inoculated intra-thoracically. (A) Mosquitoes were orally infected *via* a blood meal containing 4.10^7^ PFU/mL of YFV-DAK, YFV-17D or with no virus. Alternatively (B), mosquitoes were inoculated intra-thoracically with 2,5.10^6^ PFU of YFV-17D or YFV-DAK or with the same amount of UV-treated YFV-17D or YFV-DAK. Several whole mosquitoes were also analyzed 20 minutes after feeding or injection to ensure a similar amount of viral particles of both viral strains were delivered in mosquitoes. The relative amounts of organ-associated viral RNA were determined by RT-qPCR analysis 10 days after infection and are expressed as genome equivalents (GE) per organ. Total RNA was also extracted from several whole mosquitoes the same day of the feeding (black dots). The number of organs (n) analyzed is indicated. The dashed lines indicate the limit of detection. Data obtained from Three independent experiments were performed with untreated viruses. Control experiments with UV-treated viruses were once. Statistical analyses were performed using a Mann-Whitney test (* p < 0.05 ; ** p < 0.01 ; *** p < 0.001 ; **** p < 0.0001).

To ensure that viral RNA detected in secondary organs of injected mosquitoes represented replicative RNA and not input viral RNA, UV-treated viral RNA was also injected into the thorax of several mosquitoes. A signal, slightly above the detection threshold, was detected in two organs out of 39 tested (Fig. 4B). These data confirm the ability of YFV-17D to replicate as efficiently as YFV-DAK in midgut and secondary organs when mosquitoes were inoculated intra-thoracically.

### Genomic diversity of YFV-DAK may participate in its efficient replication and dissemination in *Aedes aegypti*

RNA viruses are comprised of a cloud of related viruses known as a quasi-species, or viral swarm [37]. This phenomenon is a direct result of the low fidelity attributed to the RNA-dependent RNA polymerase, which lacks an exonuclease proofreading function resulting in error-prone replication. This cloud of mutations may allow arboviruses to sustain a transmission cycle between insect and vertebrate hosts. Deep sequencing of YFV-17D revealed that the vaccine strain is poorly diversified compared to its parental strain Asibi [25]. This loss of genetic diversity has been proposed to contribute to YFV-17D attenuation in vaccinated patients [25]. Moreover, recent studies with Venezuelan equine encephalitis virus (VEEV), which belongs to the genus *Alphavirus*, have revealed that viruses that have disseminated in mosquitoes have an higher diversity than the ones that did not disseminate [38]. These observations prompted us to compare the diversity of the YFV-17D and YFV-DAK.

To determine a consensus sequence of the two viral strains and to analyze their variability, we performed next generation sequencing (NGS) analysis of the two viral stocks. Coverage depth for these alignments was between 200 and 300. The comparison of the two consensus sequences identified 316 synonymous mutations (Fig. 5A and 5B, blue bars) and 56 non-synonymous ones (Fig. 5B, red bars). These differences were scattered along the genome. Some of these non-synonymous mutations, as well as the 8 mutations present in the 3’ untranslated region (UTR) of the genome, could have functional consequences (Fig. 5B). Figure 5C represents the single nucleotide variants (SNVs) and their frequency all along the two genomes. Only the SNVs representing a minimum of 1% of all observations were considered. The genome of YFV-17D contained more SNVs than the one of YFV-DAK, 50 against 18. A SNV that lies in the NS2A gene of YFV-17D is represented in 44% of the population, but does not induce amino acid change. The higher abundance of variants in 17D genome as compared to DAK genome is surprising since a recent study showed that the vaccine strains YFV-17D and YFV-FNV contained fewer variants than their respective parental strains [39]. However, these observations do not inform on the general variability of the genomes since they concern only a small number of nucleotides.

**Figure 5.**
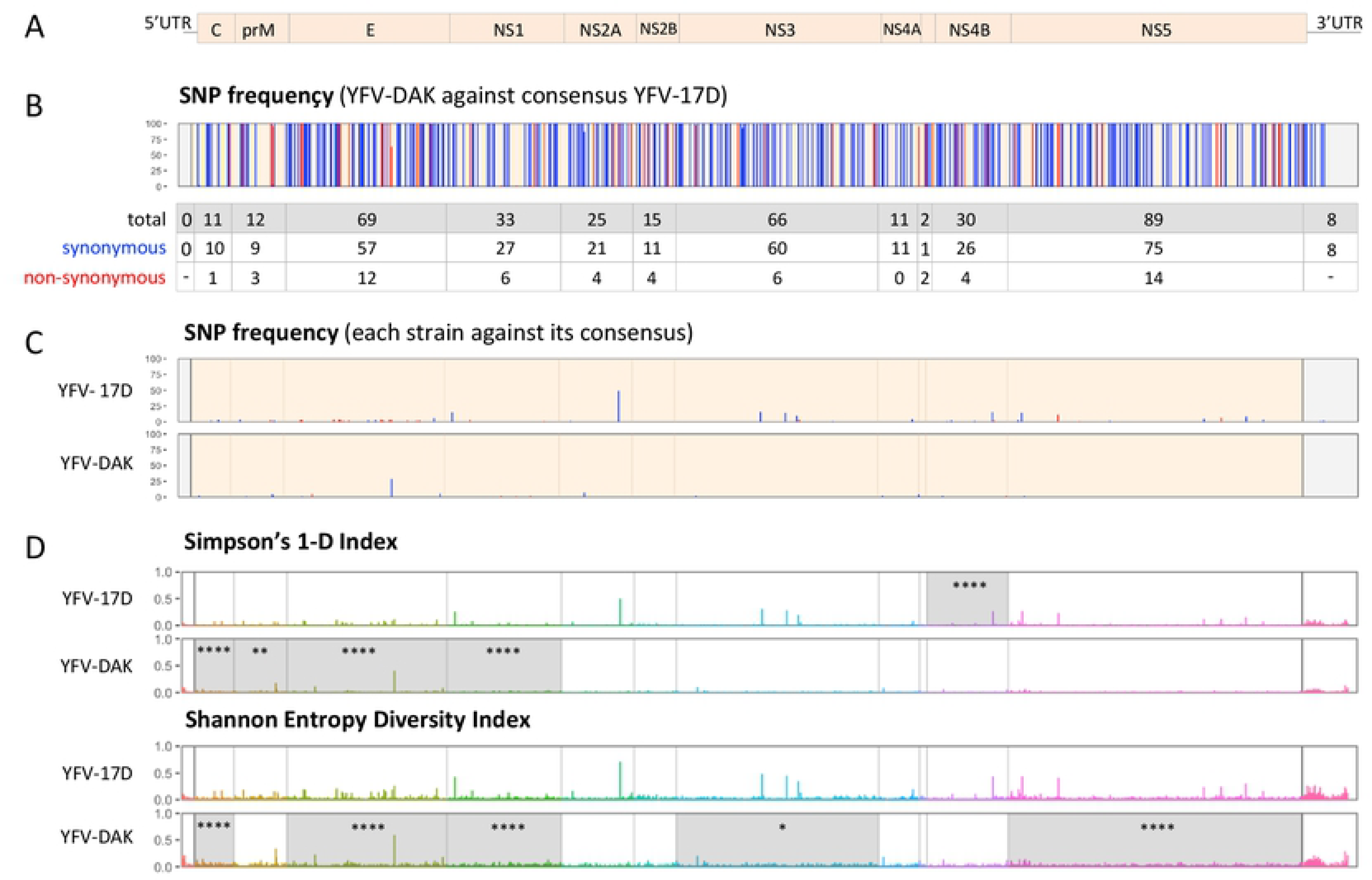
Sequencing and analysis of genome diversity of YFV-17D and YFV-DAK. (A) Schematic representation of YFV genome. (B) Genomic variability between YFV-17D and YFV-DAK. Single-nucleotide polymorphism (SNP) between YFV-Dakar and the consensus sequence of YFV-17D are shown. Blue and red bars represent synonymous and non-synonymous variants, respectively. The table shows the number of SNP per coding or non-coding region. Only SNPs of more than 1% were represented. (C) Genomic intra-variability of YFV-17D and YFV-DAK. SNP frequency were obtained using the respective consensus sequence of each strain as a reference. (D) Diversity indexes of YFV-17D and YFV-DAK are represented. Simpson’s 1-D and Shannon entropy diversity indexes were calculated and plotted for each nucleotide along the genome. Diversity indexes were compared for each gene using a Wilcoxon test (* p < 0.05 ; ** p < 0.01 ; *** p < 0.001 ; **** p < 0.0001). Genes with statistically higher diversity indexes are highlighted in grey.

We then estimated the diversity of each nucleotide of the viral genomes using the Simpson 1-D [29] and Shannon’s entropy indexes [28] (Fig. 5D). A Wilcoxon test showed that the YFV-DAK strain has a significantly greater global diversity than that of YFV-17D (Table 2) (*p* = 1.77e-03 and U = 56 957 560, *p* = 6.92e-08, and U = 56 180 896, respectively for the two indices). When comparing the diversity of each open reading frame using the two indexes, three genes of the YFV-DAK strain, coding for the C, Env and NS1 proteins, had a significantly higher diversity than their YFV-17D counterparts (dark green in Table 2). These analyzes suggest that these three proteins are major determinants of YFV-DAK diversity. As expected, when these three genes were subtracted from the analysis, the two viral genomes were no longer different in terms of genetic diversity (Table 2, bottom line). Thus, despite containing a lower number of SNVs than YFV-17D, the genome of YFV-DAK is more diverse than the one of the vaccine strain. This discrepancy between variant frequency and diversity analysis suggests that YFV-DAK variability results from a large number of mutations at a frequency below 1%. The genetic diversity of the YFV-DAK clinical strain may contribute to its ability to infect and spread efficiently in *Ae. aegypti*.

**Table 2.**
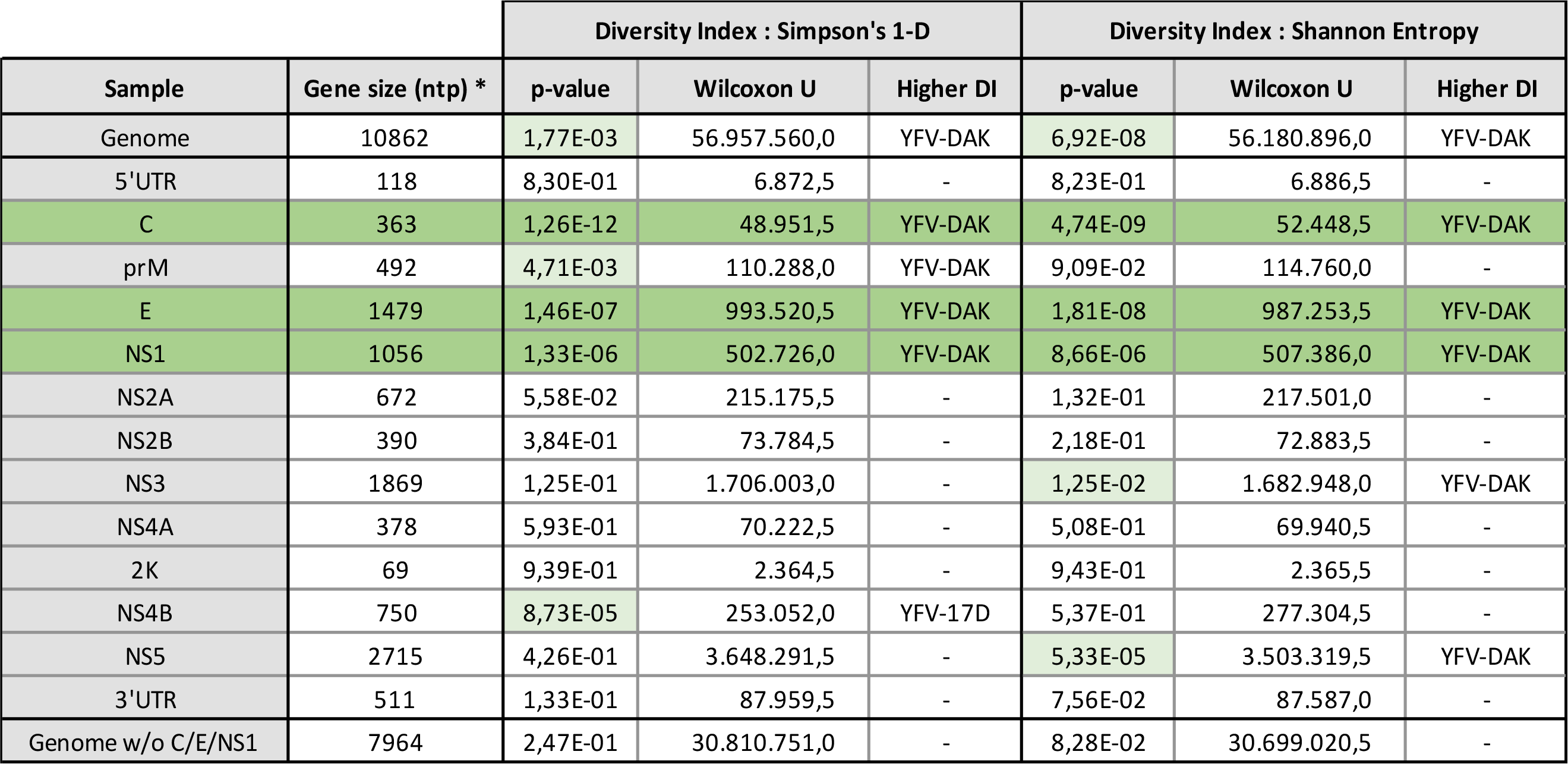
Summary table of gene diversity comparison. Statistical analysis of gene diversity using Wilcoxon test. Light green highlight genes with statistically different diversity. Dark green shows genes with statistically different genes shared between the two different indexes. * Region which are non-covered by the sequencing are not used for statistical tests. DI: Diversity Index.

### Discussion

Previous studies have shown, using titration assays on organs of the Rex-D strain of *Ae. aegypti*, that YFV-17D infects the midgut, but does not spread to secondary organs [21, 36]. Our RT-qPCR, immunofluorescence and titration analyzes revealed that the YFV-17D also replicates in the midgut of the Paea strain of *Ae. aegypti*. However, its replication was significantly less effective than the one of the clinical isolate YFV-DAK. Among the YFV-17D viruses that replicated in the midgut, less than 10% managed to reach secondary organs. None of these viruses did spread to the salivary glands. Thus, our data suggest that the YFV-17D strain is not only sensitive to the midgut escape barrier, but also to the midgut infection barrier when orally delivered. When injected into the thorax of mosquitoes, YFV-17D replicated in midgut tissues as efficiently as YFV-DAK. These data suggest that the restriction of YFV-17D replication in the midgut occur at the level of epithelial cells. Our RT-qPCR analyzes suggest that the major restriction occurs at a stage prior to viral RNA production. Several mechanisms, not mutually exclusive, could explain this restriction.

First, the restriction could occur during viral entry in midgut epithelial cells. *Flavivirus* entry mechanisms are poorly described in mosquito cells. Neither attachment factor(s) nor entry receptor(s) are identified yet. As in mammalian cells, the domain III of Env is involved in attachment and entry of the virus [40]. Thus, it is conceivable that YFV-17D Env would have a lower affinity for cell entry factors than YFV-DAK Env. Our NGS analysis revealed that the consensus sequence of the two Env proteins differs from 69 mutations, including 12 non-synonymous mutations. Seven of these non-synonymous mutations lie within the domain III. In addition, the YFV-DAK Env gene is more diverse than the YFV-17D Env gene. Finally, in Aag2 cells infected for 48 hours, we detected two forms of YFV-DAK Env under non-reducing conditions and a single form of YFV-17D Env. These differences may reflect a different conformation and may explain a different affinity for a cell entry receptor. In agreement with this hypothesis, when domain III of the Env gene of a YFV virus able to disseminate was replaced by domain III of the Env gene of YFV-17D, the dissemination of the chimeric virus was strongly inhibited, suggesting an important role in Domain III in this process [21]. These results, however, may be the consequence of chimerization, as it is known that flavivirus chimeras replicate less efficiently than parental viruses [41]. We have recently shown that the YFV-Asibi enters a panel of human cells by canonical endocytosis mechanisms involving clathrin, while YFV-17D enters cells in a clathrin-independent manner [12]. We have shown that the 12 mutations differentiating YFV-Asibi Env from YFV-17D Env are responsible for the differential internalization process. Based on these data, we hypothesized that YFV-17D and YFV-Asibi use different cell receptors [12]. It is therefore possible that the YFV-17D and YFV-DAK strains use different receptors in mosquito cells and that the receptor used by YFV-17D is poorly expressed at the apical surface of midgut epithelial cells, as compared to the one used by the clinical strain. Alternatively, the glycosylation status of the Env protein could play a role in the differential entry abilities of the two strains. Flavivirus Env proteins possess a conserved N-glycosylation motif at amino acid 153/154. This modification is involved in important viral replication and pathogenesis functions [42]. Mutagenesis studies on many flaviviruses, including the DENV, WNV and ZIKV, indicate that the loss of this N^153/154^-glycosylation impairs viral replication in the midgut [43–45]. Unlike most flaviviruses, the YFV Env lacks the N^153 / 154^-glycosylation canonical site. A second non-canonical N-glycosylation site exists at position 470. However, it is unlikely that this site is functional because it is located in the hydrophobic carboxy-terminal domain and is therefore inserted into the endoplasmic reticulum membrane. The absence of an accessible N-glycosylation site in YFV-DAK Env therefore indicates that such motif is probably not necessary for replication and dissemination in mosquitoes. We therefore believe that mutations in Env, rather than its glycosylation status, are involved in vector competence.

Another mechanism that could explain the low replication of YFV-17D in the midgut of mosquitoes is its inability to escape the antiviral mechanisms in epithelial cells. The RNA interference pathway (RNAi) is a major antiviral defense initiated by the recognition of viral replication intermediates by the Dicer-2 protein [46]. This pathway inhibits the replication of DENV and ZIKV viruses in the midgut and salivary glands of mosquitoes [47–49]. Interestingly, Myles and colleagues recently showed that the YFV C protein counteracts the RNA interference pathway in *Ae. aegypti* by protecting double-stranded viral RNA from Dicer-2-induced cleavage [50]. No amino acid sequence responsible for this effect has been identified. Our NGS analysis revealed that the consensus sequence of the C gene of our two strains of interest differ by 10 mutations, including a non-synonymous one. In addition, the YFV-DAK C gene exhibited more diversity than the one of YFV-17D. This unique mutation and/or these differences in C protein diversity could modulate their RNA interference suppression activities. It is unfortunate that Myles and colleagues [50] did not provide information on the viral strain they used.

The NS1 YFV-DAK gene is the third gene that has significantly greater diversity than its YFV-17D counterpart. Flavivirus NS1 protein is a highly conserved glycoprotein that associates as a dimer with cell membranes and is secreted into the extracellular medium as an hexamer [51]. It has recently been shown that the NS1 proteins secreted by cells infected by DENV and JEV enhance viral replication in their vectors by allowing them to escape two important antiviral mechanisms: the production of reactive species of oxygen (ROS) and the JAK / STAT pathway [52]. One can envisage that, like the NS1 proteins of DENV and JEV, YFV-DAK NS1 protein could be a potent suppressor of these two antiviral strategies. The NS1 protein of YFV-DAK could also be more expressed and/or secreted than the one of YFV-17D.

Further studies will be needed to identify the molecular mechanism (s) responsible for the low replication and dissemination of the YFV-17D vaccine strain in *Aedes* mosquito. These studies are essential to better understand the interactions between viruses and their vectors and can also contribute to the development of non-transmissible live-attenuated vaccines.

## Acknowledgments

We thank C.M Rice for generously providing the anti-YFV-NS4B antibodies; R. Kuhn for the YFV replicon construct, P. Desprès for 4G2 hybridomas and M. Flamand for anti-NS1 antibodies (17A12 and 6B8). We are grateful to the members of our laboratory for helpful discussions and technical advice. We thank Antonio Borderia for advice concerning the analysis of viral genome diversity.

